# COVID19: Exploring uncommon epitopes for a stable immune response through MHC1 binding

**DOI:** 10.1101/2020.10.14.339689

**Authors:** Folagbade Abitogun, R. Srivastava, S. Sharma, V. Komarysta, E. Akurut, N. Munir, L. Macalalad, O. Olawale, O. Owolabi, G. Abayomi, S. Debnath

## Abstract

The COVID19 pandemic has resulted in 1,092,342 deaths as of 14^th^ October 2020, indicating the urgent need for a vaccine. This study highlights novel protein sequences generated by shot gun sequencing protocols that could serve as potential antigens in the development of novel subunit vaccines and through a stringent inclusion criterion, we characterized these protein sequences and predicted their 3D structures. We found distinctly antigenic sequences from the SARS-CoV-2 that have led to identification of 4 proteins that demonstrate an advantageous binding with Human leukocyte antigen-1 molecules. Results show how previously unexplored proteins may serve as better candidates for subunit vaccine development due to their high stability and immunogenicity, reinforce by their HLA-1 binding propensities and low global binding energies. This study thus takes a unique approach towards furthering the development of vaccines by employing multiple consensus strategies involved in immuno-informatics technique.

## Introduction

Coronaviruses are a large group of viruses that are rather common throughout the community, causing illness ranging from the common cold to more severe diseases such as Severe Acute Respiratory Syndrome (SARS-CoV) and Middle East Respiratory Syndrome (MERS-CoV) (*1*). The previous outbreaks of these coronaviruses in 2003 and 2015 respectively show similarities to the novel coronavirus (*2*), which was first reported in December 2019 in Wuhan (*3*). It was later declared as a pandemic on 11th March, 2020 by the World Health Organization (WHO) having caused a global emergency across many countries and territories around the world.

The causative agent of the COVID-19 disease is the severe acute respiratory syndrome coronavirus 2 (SARS-CoV-2) which was initially tagged as 2019 novel coronavirus (2019-nCoV) by the World Health Organization (*4*) before genomic studies revealed a 79.5% and 96% nucleotide level similarity with SARS-CoV and bat coronavirus respectively (*5,6*), hence necessitating its new identity. Further analysis on its similarity with SARS-CoV and MERS-CoV including phylogeny revealed that it is categorized under the family Coronaviridae, order Nidoviralae and genus Betacoronavirus alongside the other two viruses which have also caused pandemic situations in the past, causing thousands of death globally. (*7,8*)

The structural architecture of SARS-CoV-2 is an elaborate one, containing a positive single stranded RNA as its genetic element with genomic length of about 29-30 kilobases, (*9*) which encodes the major structural proteins being the spike (S) glycoprotein, membrane (M) protein, envelope (E) protein, and nucleocapsid (N) protein. (*10,11*) The structure of the spike glycoprotein of the virus is also an extended similarity with SARS-CoV, (*4*) which together with COVID19: Exploring uncommon epitopes for a stable immune response through MHC1 binding other proteins of the virus are candidates for vaccine development and are being explored in different settings due to the active roles of the proteins in the infectivity of the virus. (*12–15*) Developing an effective treatment for SARS-CoV-2 is a research priority. Currently, different therapeutic strategies are being utilized by researchers to combat this dangerous COVID-19, with much of the focus being on developing novel drugs or vaccines. However, in recent years, the development of vaccine design has been revolutionized by the reverse vaccinology (RV), which aims to first identify promising vaccine candidate through bioinformatics analysis of the pathogen genome. One of the successes of this method is the Bexsero vaccine, discovered for Group B meningococcus. (*16*) This immuno-informatics method includes the identification of peptide epitopes in the virus genome which can then be prepared by chemical synthesis techniques. These epitopes are easier in quality control; however, there is the need for structural modifications as well as the inclusion of delivery systems, and adjuvants in their formulation due to the low immunogenicity of the epitopes which is a result of their structural complexity and low molecular weight. (*17*)

Recently, a set of B and T cell epitopes highly conserved in SARS-CoV-2 were identified from the S and N proteins of SARS-CoV. (*18*) However studies have shown that full length spike protein vaccines for SARS-CoV may lead to antibody mediated disease enhancement causing inflammatory and liver damage in animal models (*19,20*) which is why in this manuscript, we applied immuno-informatics “in silico” approaches to identify potential CD8+ cytotoxic T Cell epitopes from proteins of SARS-CoV-2, SARS-CoV and MERS-CoV. In achieving this, the ~29.8kb genome of the SARS-CoV-2 which encodes for 28 proteins comprising 5 structural proteins, 8 accessory proteins and 15 non-structural proteins as well as the ~30kb genome of SARS-CoV-1 comprising 14 open reading frames (ORF) were explored for some uncommon proteins of the viruses with therapeutic potentials. Identified proteins were thus considered for highly antigenic epitope prediction and evaluation. The antigenicity and immunogenicity of all identified epitopes was estimated and their interactions with the human leukocyte antigen (HLA) class I system were evaluated, owing to the wide usage of the HLA system as a strategy in the search for the etiology of infectious diseases and autoimmune disorders, (*21*) as was explored in the case of SARS-CoV-1 after the first epidemic that broke out in East Asia in 2002-2003. This is linked to the fact that HLA genes are known to display the highest level of diversity in the genome, with thousands of different alleles now reported. Each of these alleles is also a combination of multiple single nucleotide polymorphisms (SNPs). This, we also investigated the HLA-class I system for evidence of disease association, with the hope that the discovery of disease susceptibility will help in developing protection such as vaccines against the virus.

## Methods

### Protein sequence retrieval

The UniProt (*22*) database was searched for the protein sequences inferred from the genome sequence analysis of different isolates of highly pathogenic human coronaviruses. The SARS CoV, MERS-CoV, and SARS-CoV-2 proteins without experimentally defined 3D structure or functional annotation were chosen for further analysis and their sequence files in the FASTA format were downloaded.

### Sequence analysis and physicochemical parameters evaluation

The FASTA sequences were used to infer the physicochemical properties of the chosen uncharacterized viral proteins. With tools and servers including ExPASyProtParam, (*23*) EMBOSS Pepstats (*24*), Peptide 2.0, (*25*) PROTEIN CALCULATOR v3.4, (*26*) important physicochemical properties of the proteins including the sequence length and molecular weight, theoretical isoelectric point (IpH), hydrophobicity/hydrophilicity and polarity indices (calculated as the percentage of hydrophobic, hydrophilic and polar amino acids respectively), the content of the basic amino acids (arginine, histidine, lysine), and disulfide bond allowing cysteine were evaluated and computed. No less than three online tools were used to calculate each parameter; the consensus results were chosen for presentation in this paper.

### Tertiary structure modeling

The 3D structures of the retrieved viral proteins were obtained using different homology modeling tools of which at least four are approved by CASP. (*27*) Homology 3D protein modeling against the protein templates with experimentally defined tertiary structure was done using the following online servers: SWISS-MODEL, (28) LOMETS, (29) IntFold, (*30*) I-TASSER, (*31*) FALCON, (*32*) GalaxyWeb. (*33*) The other tools used utilize different approaches: RaptorX (*34*) exploits distance-based protein folding powered by deep learning; QUARK (*35*) implements ab initio protein folding based on building from small fragments by replica-exchange Monte Carlo simulation under the guide of an atomic-level knowledge-based force field. Application of different algorithms lying behind each modeling tool provided various variants of 3D structure models for each protein. Up to five top models of each protein resulting from every tool were selected for further processing.

### Refinement and validation of tertiary structure

Each model was refined using a molecular dynamics simulation approach implemented by GALAXY Refine (*36*) tool or rapid energy minimization using discrete molecular dynamics with an all-atom representation for each residue implemented by Chiron server. (*37*) The top refined model was chosen for every protein based on the refinement servers output model quality indices. The further Ramachandran analysis with PROCHECK (*38*) and Saves v5.0 tools Verify3D, (*39*) ERRAT, (*40*) and PROVE (*41*) was performed on the top models of all proteins initially included in the analysis. Thermodynamic stability of the molecules was predicted using SCooP analysis. (*42*) GROMOS96 (43) force field calculations were applied to identify the most stable protein structures. Eight (8) protein models with the most geometrically and thermodynamically stable structures that most likely represent the native 3D structures of coronavirus proteins were finally selected.

### Antigenicity prediction

Antigenicity prediction of the top eight uncharacterized coronavirus proteins selected after 3D molecular models’ validation was performed using VaxiJen tool. (*44*) A threshold value of 0.4 was taken into account. Non-antigenic peptides having VaxiJen scores less than 0.4 were discarded, while antigenic epitopes with scores higher than 0.4 were further prioritized for their immunogenicity. The proteins predicted to be non-antigens were discarded from the further analysis.

### T Cell Epitope Prediction and binding MHC class I alleles

Antiviral immunity relies on the ability of major histocompatibility complex (MHC) class I molecules to bind antigen molecules and display them to T cells. In humans, MHC class I is represented by human leukocyte antigens HLA-A, HLA-B, and HLA-C that are very polymorphic among human populations. The most common for African and Asian populations HLA-A, HLA-B, and HLA-C subtypes were retrieved from Allele Frequency Net Database. (*45*) NetMHCpan 4.1 server (*46*) which applies artificial neural networks (ANNs) trained on a combination of more than 850,000 quantitative Binding Affinity (BA) and Mass-Spectrometry Eluted Ligands (EL) peptides to predict the binding of peptides to any MHC molecule of known sequence was used to predict strong binding epitope sequences from these top 10 models. The strong binding epitope sequences predicted from the NETMHCpan server were then analyzed for their antigenicity using Vaxijen (*45*) (threshold = 0.4). Further analysis including immunogenicity, allergenicity and toxicity of the peptide epitopes were done using IEDB Immunogenicity analysis resource (Class I Immunogenicity), (*47*) AllerTOP v. 2.0 (*48*) and ToxinPred (*49*) servers respectively. Peptide epitopes with the most immunogenic sequences which were non allergens and nontoxic were used for molecular docking simulations with specific HLA-1 alleles.

### Molecular docking between MHC class I and predicted peptide epitopes 3D structures

The 3D structures for the most immunogenic sequences were generated using PEPFOLD3 server (*50*) which applies a de novo approach based on a new Hidden Markov Model sub-optimal conformation sampling to predict models for peptides from 5 to 50 amino acids. HLA structures based on the binding predictions from NETMHCpan were downloaded from RCSB PDB and isolated from their complexes using Chimera. (*51*) Molecular docking of these molecules was carried out by PatchDock (*52*) which employs a surface patch matching algorithm inspired by object recognition and image segmentation techniques to dock the molecules. The top 1000 results from PatchDock were then submitted to FireDock (*53*) for refinement of protein-protein docking solutions. Given a set of up to 1000 potential docking candidates, FireDock refines and scores them according to an energy function, spending about 3.5 seconds per candidate solution. Each candidate is subsequently refined by restricted interface side-chain rearrangement and by soft rigid-body optimization. The side-chain flexibility is modeled by rotamers and the obtained combinatorial optimization problem is solved by integer linear programming. Following rearrangement of the side-chains, the relative position of the docking partners is refined by Monte Carlo minimization of the binding score function. The refined candidates are ranked by the binding score which is a combination of Atomic Contact Energy, softened Van der Waals interactions, partial electrostatics and additional estimations of the binding free energy.

### Conservancy Analysis

In order to identify functionally important regions in the selected proteins and the selection of epitopes with the desired degree of conservation, Consurf (*54*) was used to determine the variability of epitopes within the selected proteins. The degree of conservation at each amino acid site is similar to the inverse of the site’s rate of evolution; slowly evolving sites are evolutionarily conserved while fast evolving sites are variable. (*55*) Amino acid residues that are crucial for retention of protein function are believed to be associatedwith inherently lower variability, even under immune pressure and as such often represent good targets for the development of epitope-based vaccines. (*56*) Both 3D structure and FASTA sequences of the proteins were used as inputs and the Consurf server carries out both a PSI-BLAST search for close homologues and a multiple sequence alignment using CLUSTAL W. The server thus builds a phylogenetic tree that is consistent with the MSA and then calculates the conservation grades which are color-coded, taking into account the evolutionary relationship between the homologues.

## RESULTS

### Human Coronavirus uncharacterized protein sequences analysis

A total of 8 non-structural sequences were identified and retrieved from UniProtKB database. The mass, isoelectric-pH, Basic amino acid %, Cysteine %, Polarity and Hydrophobicity of the protein sequences were computed and outlined in table 1. The lengths of these protein sequences varied from 39 to 275 amino acid residues. Cysteine was found to have the highest presence in the selected proteins with an average of 5.56% and its highest presence was observed in proteinQ7TFA0 with 15.40%. Arginine, Lysine and Histidine have an average presence of 2.35%, 3.69% and 4.36% respectively. The polarity and non polarity of the final selected proteins ranged from 27.907%to 63.839 % and from 36.161%to 72.093% respectively with M4SVF1 having the highest percentage of polar amino acids (63.839%) while P0DTD8 has the highest percentage of non-polar amino acids (72.093%). Likewise, the highest hydrophobicity index was recorded for P0DTD8.

**Table 1.**
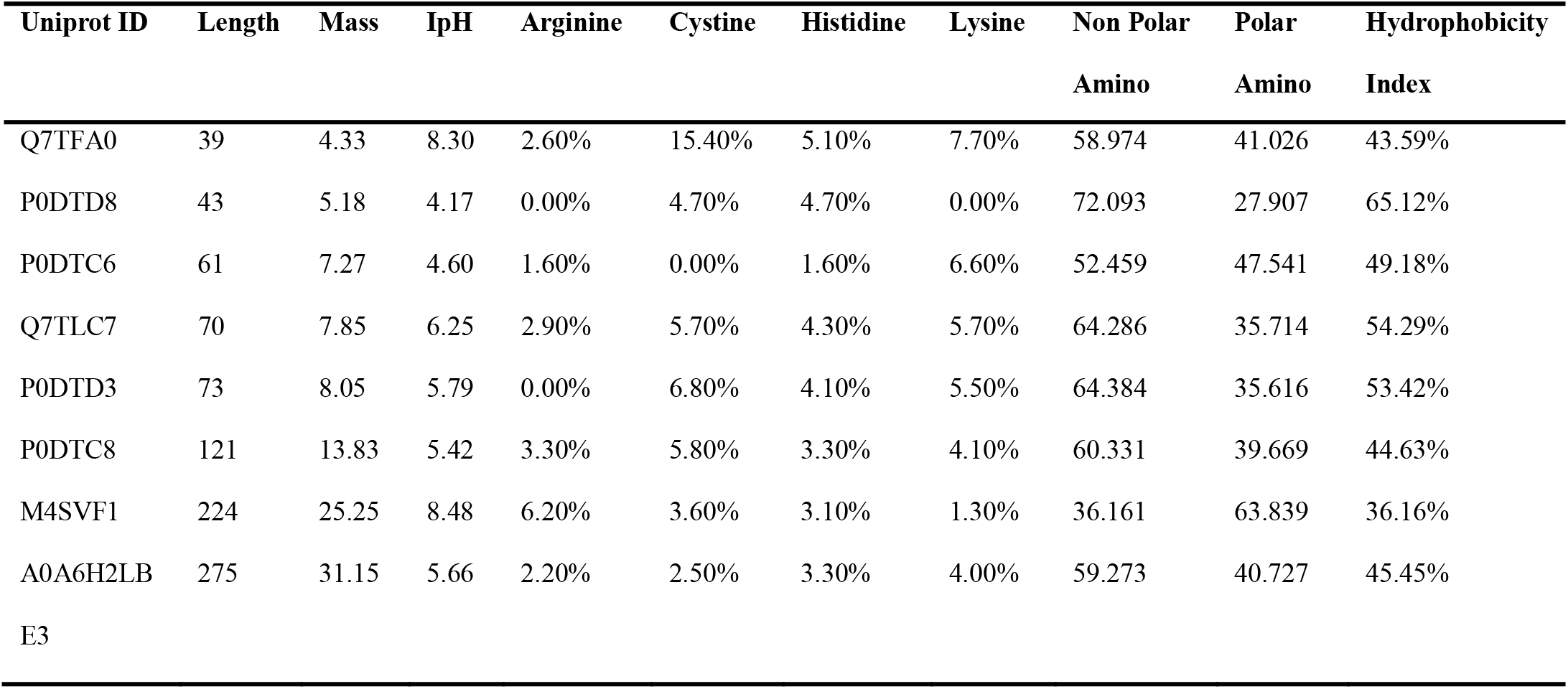
Analysis of Identified Protein Sequences

### Protein 3D Structure Predictions

After obtaining the 3D structures of the proteins from modeling servers, the models were refined using GalaxyRefine. Upon refinement, the models were subjected to stereochemical and thermodynamic stability analysis. Particularly, the models with highest Ramachandran favored residues and lowest clash scores were used for energy analysis using Chiron, SCooP and ERRAT servers as highlighted in table 2. All the protein structures had a Ramachandran score of over 89% and an ERRAT value of over 60, with Q7TFA0, P0DTD8 and P0DTD3 all having a perfect 100% Ramachandran scores. The free energy of the structures ranged from −28.7kcal/mol (Q7TFA0) to −2.2 kcal/mol (A0A6H2LBE3). Force field values ranging from −5955.989 (A0A6H2LBE3) to −435.783 (P0DTD8) were observed using GROMOS96 implementation of the Swiss-PDB Viewer. The models with lowest overall energies were determined to be stable.

**Table 2.** Stability Analysis of the predicted 3D structures.

| UniProt ID | ERRAT Values | GROMOS96 Force field values | $\Delta G_r$ kcal/mol | RC Aanalysis | Tm °C |
| --- | --- | --- | --- | --- | --- |
| Q7TFA0 | 94.1176 | -549.952 | -28.7 | 100.00% | 114.3 |
| Q7TLC7 | 93.5484 | -2728.715 | -9.6 | 89.30% | 65 |
| P0DTD3 | 84.9057 | -1550.547 | -11.3 | 100.00% | 69.4 |
| M4SVF1 | 84.7222 | -6842.415 | -5.7 | 92.20% | 70.9 |
| P0DTC6 | 83.0189 | -1689.699 | -13.7 | 94.80% | 85.5 |
| P0DTD8 | 80.6452 | -435.783 | -20.7 | 100.00% | 101.7 |
| P0DTC8 | 71.4286 | -1854.744 | -4.3 | 98.10% | 84.3 |
| A0A6H2LBE3 | 64.557 | -5955.949 | -2.2 | 95.20% | 73.6 |

### Viral Protein Antigenicity Predictions

VaxiJen Antigenicity prediction was used to predict the overall antigenicity of the 8 proteins as outlined in table 3. Two proteins: M4SVF1_MERS and Q7TFA0 were predicted to be non-antigens with their antigenicity scores of 0.3373 and 0.1251 respectively while P0DTD8 and P0DTC8 showed very high antigenicity owing to their high scores of 0.8462 and 0.6502 respectively.

**Table 3.**
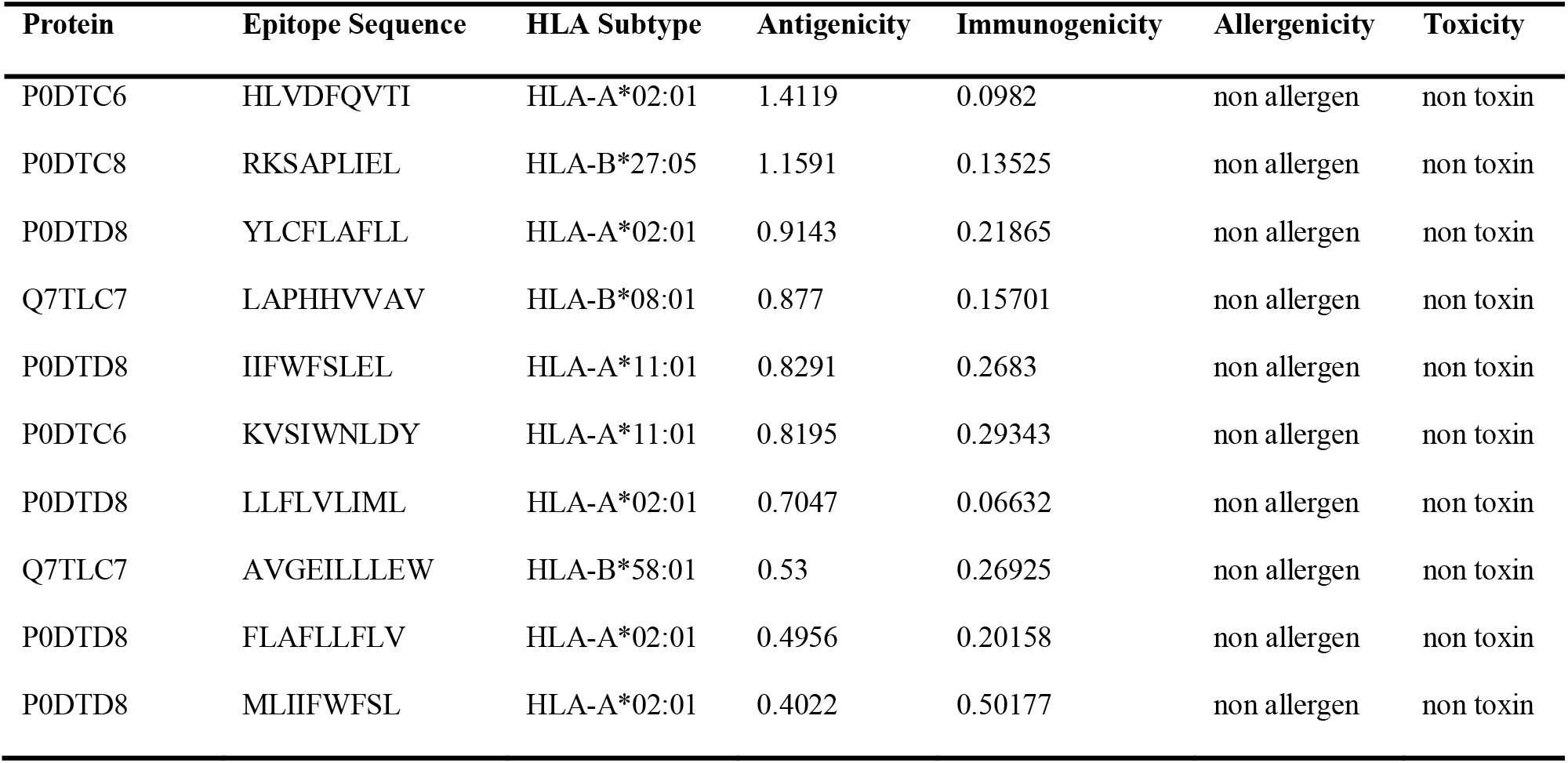
Immunogenicity analysis of predicted MHC Class 1 epitopes.

### Human Leukocyte Antigen Interactions

T-cell epitope prediction was carried out using IEDB, EpiJen, PepVac, Rankpep, and NetMHC. Initially, 15,181 HLA class I binding epitopes were predicted from 6 out of 8 stable proteins which were predicted to be antigenic. Scrutiny on the basis of percentile rank filtered 112 peptide epitopes. Considerable binding affinity for 5 HLA-1 subtypes with high frequencies in Asia and Africa (HLA-A*02:01, HLA-A*11:01, HLA-B*08:01, HLA-B*27:05 and HLA B*58:01) was observed from NETMHCpan server which predicted high affinity binders (<0.5 percentile rank) using EL-Ranks. Among predicted epitopes of SARS-CoV-2 virus, ten (10) epitopes showed considerably high immunogenicity towards the MHC-Class 1 molecules which were then selected for further analysis (Table 4). All the predicted epitopes were further subjected to allergenic and toxicity analysis, with all 10 coming out as non-allergens and non toxins (Table 4). Epitopes including HLVDFQVTI, RKSAPLIEL, and YLCFLAFLL showed remarkably high antigenic tendencies, scoring 1.4119, 1.1591 and 0.9143 respectively. All of these epitopes, along with their features are reported in Table 4. The structures of HLA-1 Subtypes namely, HLA-A*02:01, HLA-A*11:01, HLA-B*08:01, HLA-B*27:05 and HLA-B*58:01 were obtained from RCSB PDB (Table 5). Further docking analysis resulted in a total of 10 global binding energies predictions which were refined with FireDock (Table 6). The global binding energies range from −68.71 to −11.19 indicating that all the proposed epitope sites from the modelled proteins can serve as good antigens.

**Table 4.**
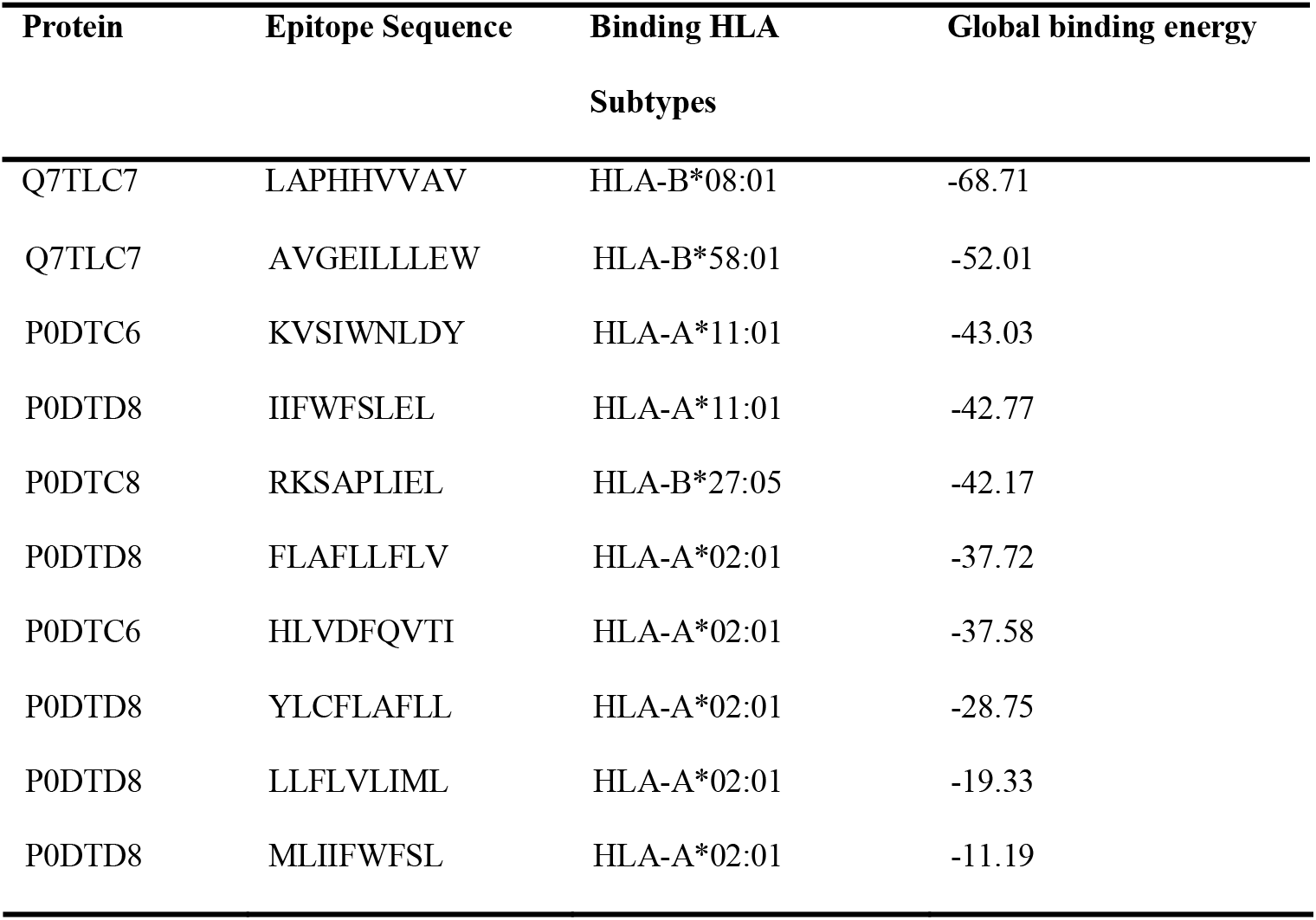
Global binding energies of the predicted epitope sequences

**Table 5.** PDB ID of selected HLA Subtypes

| HLA Subtype | PDB ID | Name |
| --- | --- | --- |
| HLA-A*02:01 | 4U6X | Crystal Structure of HLA-A*0201 in complex with ALQDA, a 15 mer self-peptide |
| HLA-A*11:01 | 5WJN | Crystal Structure of HLA-A*11:01-GTS3 |
| HLA-B*08:01 | 4QRQ | Crystal Structure of HLA B*0801 in complex with HSKKKCDEL |
| HLA-B*27:05 | 2BSS | Crystal structures and KIR3DL1 recognition of three immunodominant viral peptides complexed to HLA-B2705 |
| HLA-B*58:01 | 5INC | Crystal structure of HLA-B5801, a protective HLA allele for HIV-1 infection |

### Conservation Analysis of the Selected T-cell epitopes

The conservation status of each residue in the selected T-cell epitopes in four (4) SARS COV-2 proteins (Q7TLC7, P0DTC6, P0DTD8, P0DTC8) were examined using ConSurf. Results revealed that the highly conserved and exposed residues are found in all 4 proteins, with P0TDC6 and P0DTC8 having the larger amount of these residues. Particularly, the epitopes without allergenicity and toxicity containing at least one predicted functional residue (highly conserved and exposed) included epitopes ‘RKSAPLIEL’ and ‘HLVDFQVTI’ while T-cell epitopes ‘RKSAPLIEL’, ‘FLAFLLFLV’, and ‘HLVDFQVTI’ contained at least one predicted structural residue (highly conserved and buried). Interestingly, none of the epitopes from P0DTD8 contained any functional residues (highly conserved and exposed) while the highest concentration of predicted structural and functional residues was found in two epitopes HLVDFQVTI (1 functional residue and 3 structural residues) and RKSAPLIEL (3 functional residues and 1 structural residue)-from the protein P0DTC6.

## Discussion

Despite the continuous unrest caused by the COVID19 pandemic, considering the 1,886,643 new cases recorded as of 7th September, 2020 which has spiked up the total reported cases to over 26.7 million and a total of 876,616 deaths globally, there is currently no available FDA-approved vaccine against COVID-19 (*57*). If a vaccine is successfully developed against COVID-19, this will improve global human health. Advancement in Technology has led to various Bioinformatic approaches such as Reverse Vaccinonology (RV) and immune-informatics being applied to revolutionize vaccine production in terms of production cost and time which has been a major setback in vaccine development. Antibody response as well as cell mediated immunity can be established by using proper protein antigen. Reverse vaccinology (RV) has been useful in identification of potential vaccine candidates against pathogens and the world is in a race to get one against SARS-COV2. This approach presents an advantage over other vaccine approaches because it can identify all potential antigens coded by a genome irrespective of their abundance, phase of expression and Immunogenecity.

While most of the previous protein vaccines against SARS-CoV have focused on the full length spike protein of the virus, the vaccine development race against the COVID19 virus have followed the same path, with a larger percentage of the 69 of the 210 vaccines currently in development utilizing the spike protein of the SARS-CoV-2 (*58*) not minding the reported negative side effects, including liver damage and antibody mediated inflammatory diseases, of the full length spike protein vaccines and other whole organism vaccines in the host. Additionally, while novel technologies and approaches such as Next-generation vaccines enabled through advances in nanotechnology which relies on the principle that nanoparticles and viruses operate at the same length scale are poised to make a clinical impact for the first time, many of these platforms may be several years away from deployment and therefore may not have an impact on the current SARS-CoV-2 pandemic, further necessitating the priority given to the quicker and safer vaccine development platforms. The current study focused on the HLA-1 binding propensities of some uncommon proteins from COVID-19 virus. Obtaining these antigenic epitopes using the immune-informatics approach will help inform which epitopes to use in the construct of protein vaccines against SARS-CoV-2 using proteins other than the spike protein. From the 8 selected proteins obtained all the models with good stability analysis including the Ramachandran analysis, which is a two dimensional plot drawn in the space of angles ϕ and ψ that quantify the rotation of the protein backbone and thus play a crucial role in the secondary structure of proteins and later the tertiary structure, were further analyzed for antigenicity and only highly antigenic proteins were selected for further analysis since the highly antigenic proteins will induce better immune response in the host.

Five (5) of the 35 most common HLA-1 subtypes in Africa and Asia that were identified in this study were included in the analyses and their potential interactions between identified epitopes from the proteins were predicted using the NetMHCpan 4.1 server. The binding affinities between these HLA subtypes and the epitopes were indicated by the server which allowed the selection of epitopes that will have strong interactions with the HLA alleles. The analysis of this interaction is important because vaccination is a proven strategy for protection from disease and an ideal vaccine would include antigens that elicit a safe and effective protective immune response. HLA-restricted epitope vaccines, including T-lymphocyte epitopes restricted by HLA alleles, embodies a newly developing and favorable immunization approach. Research in HLA restricted epitope vaccines to be used for the treatment of tumors as well as for the prevention of viral, bacterial, and parasitic infections have, in recent years, achieved substantial progress and this further strengthens the resolve of the HLA-binding propensities analysis done in this study. The predicted strongest binding epitopes were thus further subjected to immunogencity evaluation which checked the ability of the epitope to induce an immune response and also predicted how strongly they will induce an immune response; Allergenicity which verified the potential of our epitopes to cause sensitization or allergic reactions related to IgE antibody responses; and Toxicity which is its ability to cause bodily harm. Using these criteria, 10 epitopes were identified and these could be potential epitopes for vaccine constructs against SARS-CoV-2. This was also evinced by the lower global binding energies between our identified epitopes and the HLA molecules, ranging from −68.71 to −11.19, indicating stable complexes formed between them.

## Conclusion

Our extensive approach towards the prediction of stable protein structures from previously uncharacterized proteins and the subsequent application of immune-informatics for the identification of immunogens has resulted in the classification of 10 different epitope sites from 8 different proteins that have high antigenicity and low binding energies with HLA-1 alleles that are most commonly present in the continent of Asia and Africa. These epitopes have high tendencies to provide effective and strong protective cytotoxic immunity against the SARS CoV-2 if they are adequately exploited for vaccine production. A multiple consensus approach ensures the reliability and reproducibility of these results. This study thus provides further knowledge on the association of these HLA-1 alleles with COVID19, suggesting that good peptide coverage can be achieved by many different combinations of HLA-1 alleles.

**Figure 1:**
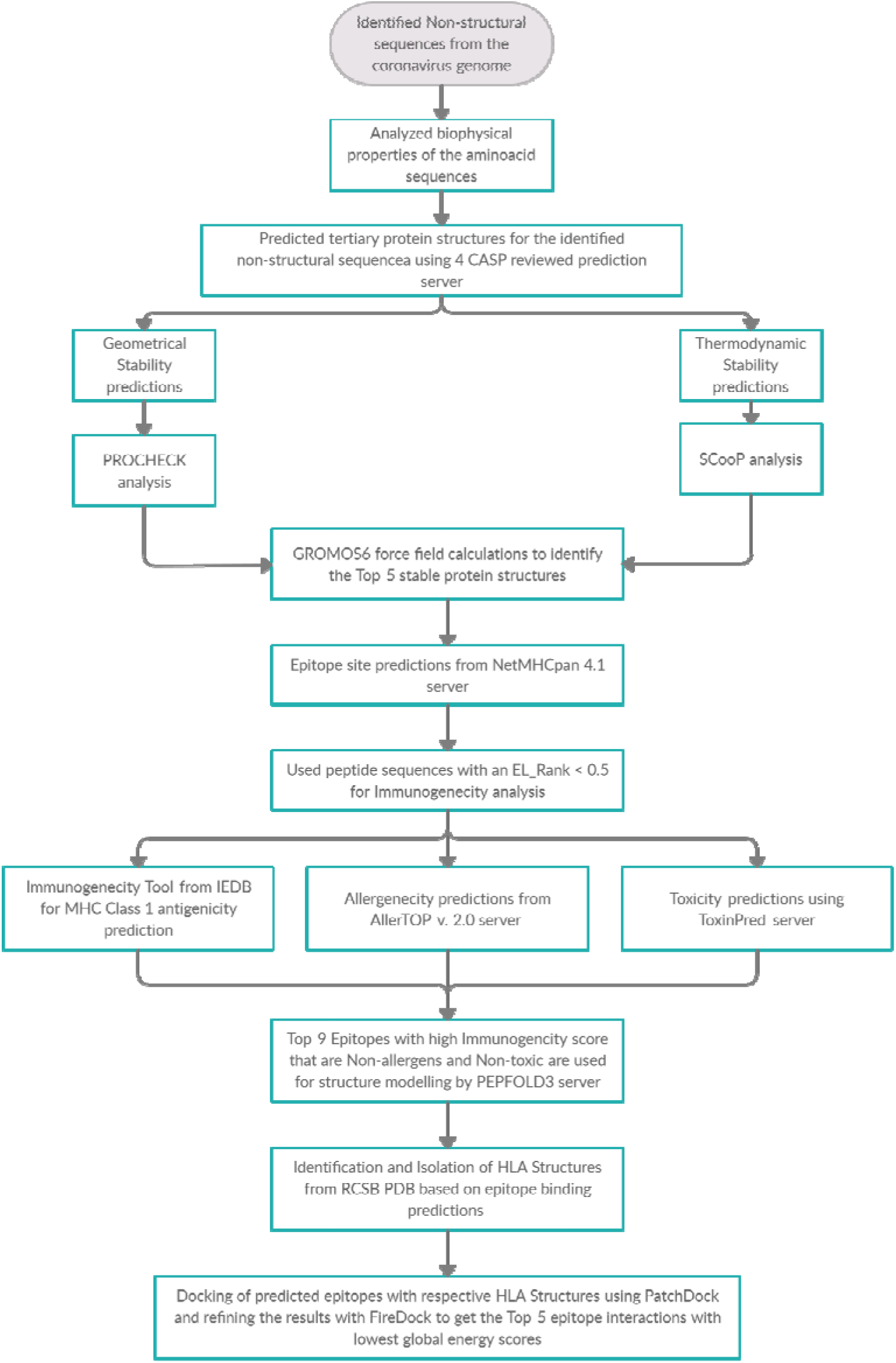
Schematic representation of the workflow:

**Figure 2:**
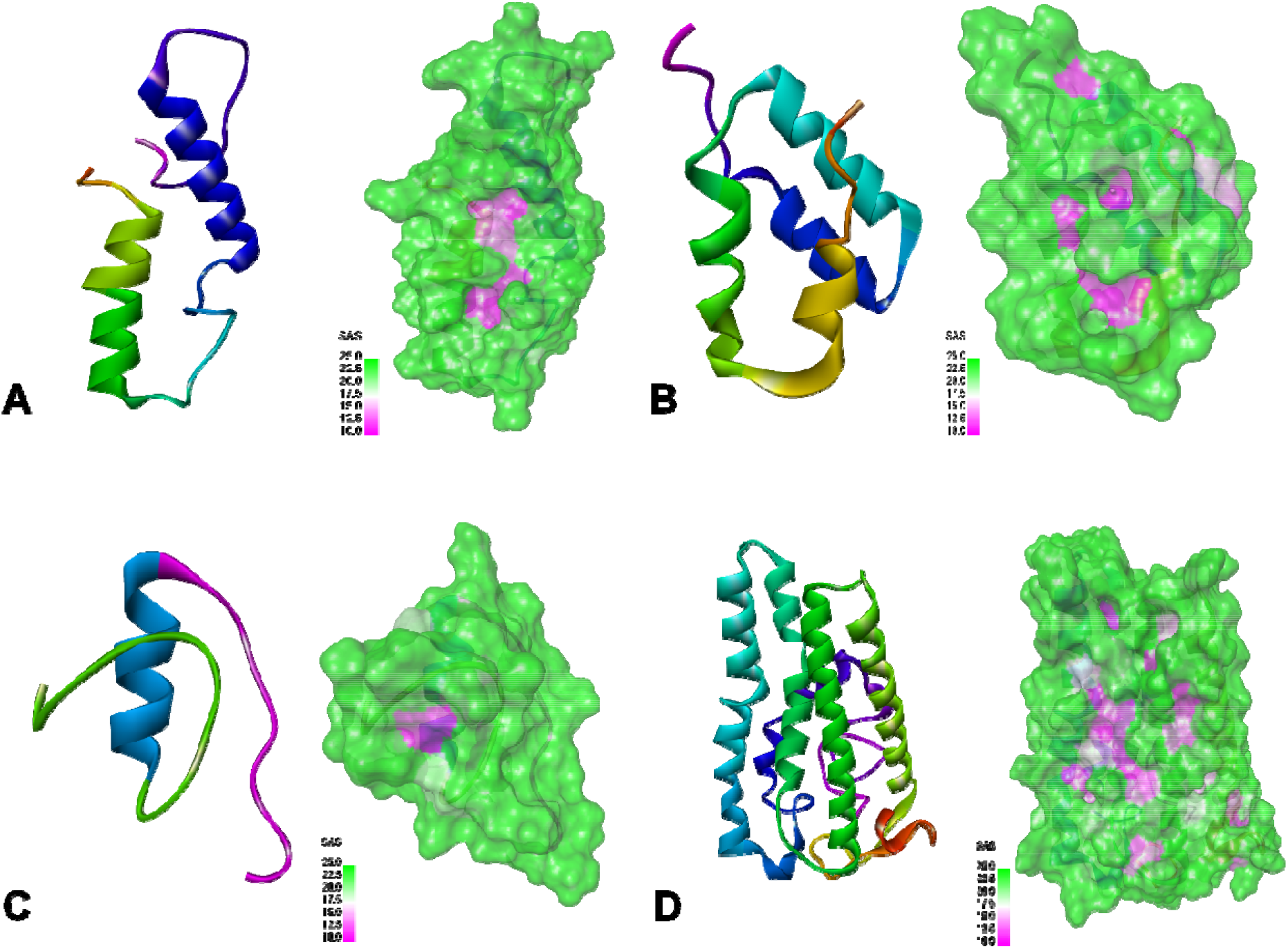
The predicted structures of the top 4 most stable proteins with their solvent accessibility surface diagrams. The predicted models were visualized using Biovia Discovery Studio Visualizer (A) Q7TFA0 (B) Q7TLC7 (C) P0DTD3 and (D) M4SVF1.

**Figure 3:**
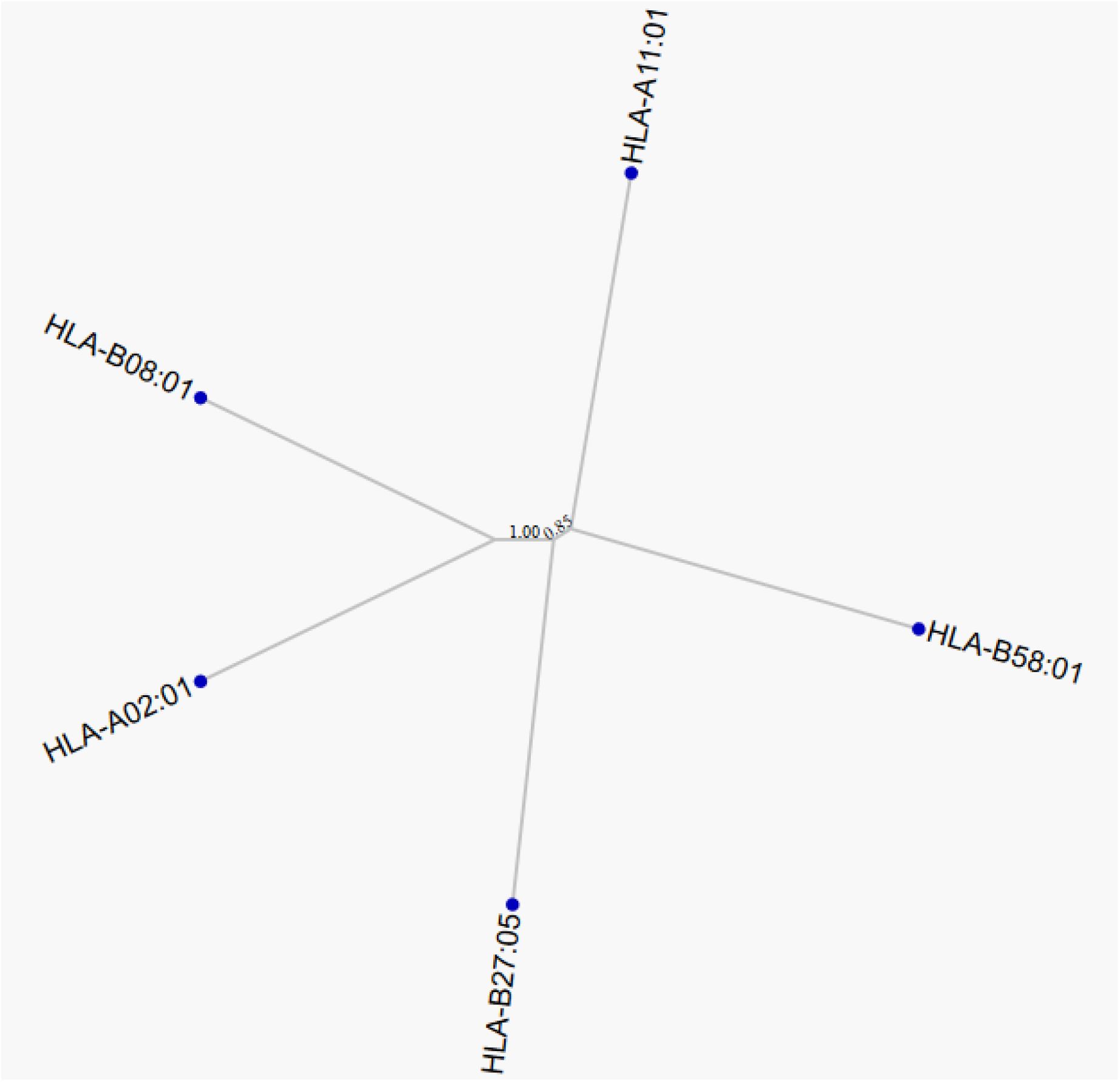
Phylogenetic tree of the 5 HLA Subtypes analyzed in this study generated using MHC-Cluster 2.0 Server^60^

**Figure 4:**
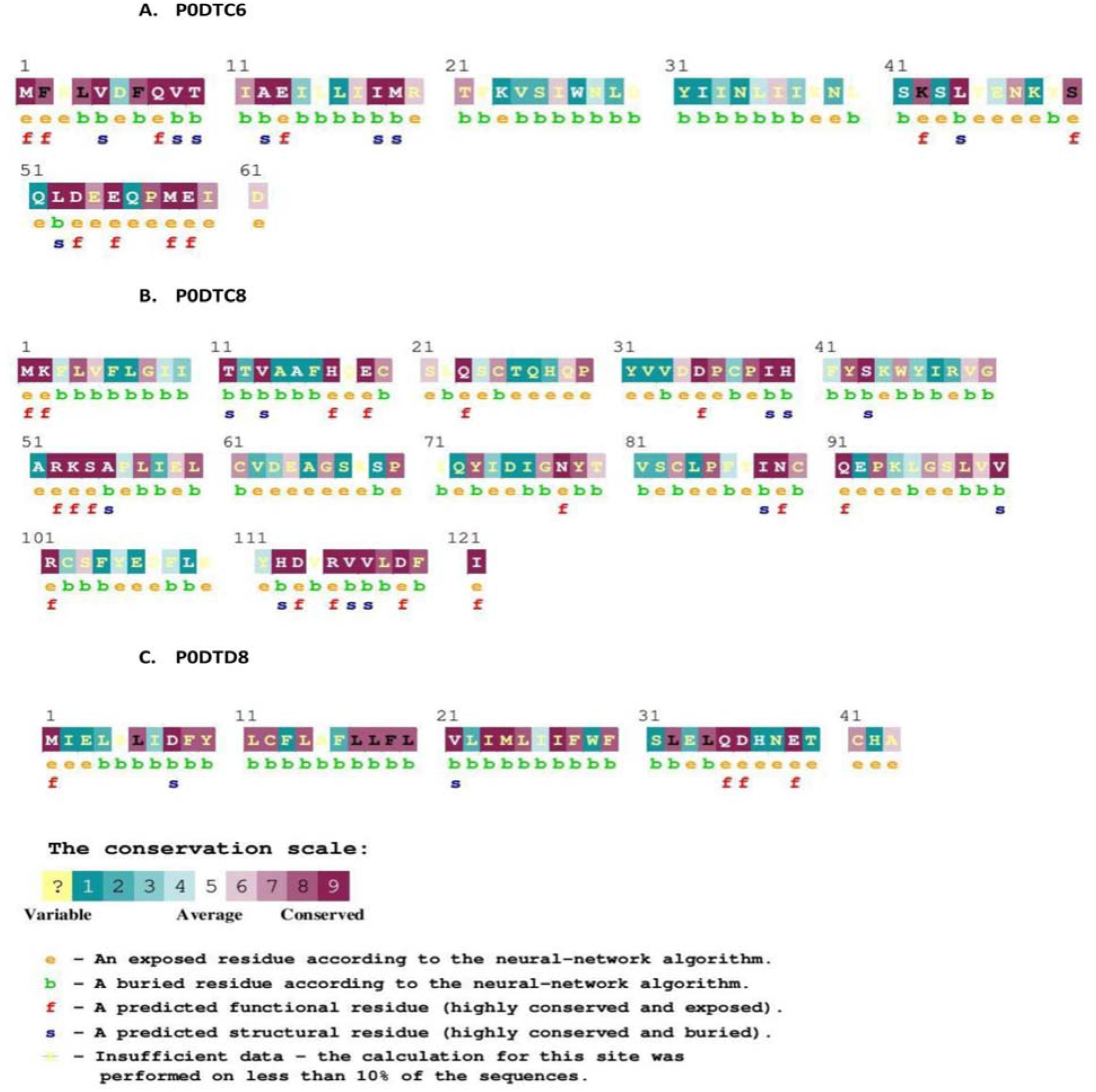
Conservation of T-Cell Epitopes in SARS-CoV-2. The position of conserved epitopes in the protein sequence. (A) P0DTC6 (B) P0DTC8 (C) P0DTD8. According to the neural network algorithm, The orange coloured ‘e’ represents exposed residues, the green colored ‘b’ represents buried residues, the red colored ‘f’ represents functional residues (highly conserved and exposed) and the dark blue colored ‘s’ represents structural residues (highly conserved and buried). The conservation scale represents the status of conservation from variable, average to conserved.

**Figure 5:**
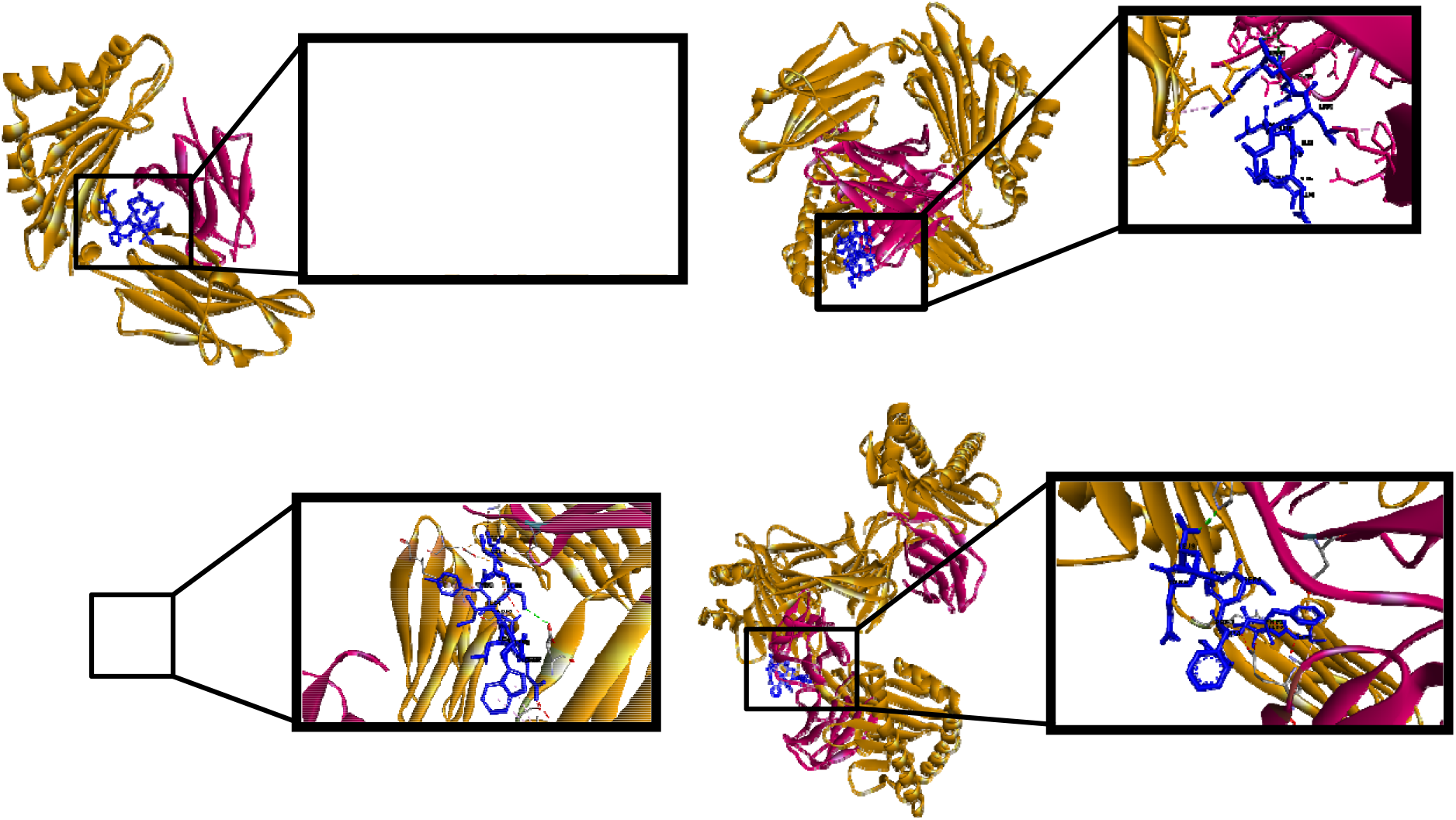
Graphical presentation of the predicted interactions between MHC class 1 binding T cell epitopes from the modelled proteins and HLA alleles. (A) Epitope: LAPHHVVAV bound to HLA-B*08:01 with a global binding energy of −68.71. (B) Epitope: AVGEILLLEW bound to HLA-B*58:01 with a global binding energy of −52.01. (C) Epitope: KVSIWNLDY bound to HLA-A*11:01 with a global binding energy of −43.03. (D) Epitope: IIFWFSLEL bound to HLA-A*11:01 with a global binding energy of −42.77.

## Acknowledgments

We acknowledge the support of DNACompass, CMESBAHF Nigeria, BIOTRUST Scientific, Galaxy Project and NeliRef for this project. We also acknowledge the provision of figure images generated by Rajavee Srivastava and Folagbade Abitogun from UCSF Chimera, Discovery Studio, PyMol, MHC Cluster Server v 2.0 and Consurf. We acknowledge T. Thao, H. Aminu, and V. Ekundayo for their contribution to this project.

## Author Bio

Folagbade Abitogun is a Research Associate at the University College Hospital, Ibadan, Nigeria. He is currently actively involved in antimicrobial resistant research and he has deep interest in the interplay between the human immune system and the clearance of infectious diseases, using molecular and bioinformatics approaches.

